# Net reductions in greenhouse gas emissions from feed additive use in California dairy cattle

**DOI:** 10.1101/2020.05.26.116277

**Authors:** Xiaoyu Feng, Ermias Kebreab

**Affiliations:** Department of Animal Science, University of California, Davis, California, United States of America

**Author notes:** Corresponding author, (EK). Author Contributions: Conception, designation, methodology: EK XF Data collection, formal analysis, model development, data analysis, original draft writing: XF Funding acquisition, project administration, final review and editing: EK.

**Keywords:** life cycle assessment, dairy cow, methane, feed additive

## Abstract

The livestock industry is one of the main contributors to greenhouse gas emissions and there is an increasing demand for the industry to reduce its carbon footprint. Several studies have shown that feed additives 3-nitroxypropanol and nitrate to be effective in reducing enteric methane emissions. The objective of this study was to estimate the net mitigating effect of using 3-nitroxypropanol and nitrate on total greenhouse gas emissions in California dairy industry. A life cycle assessment approach was used to conduct a cradle-to-farm gate environmental impact analysis based on dairy production system in California. Emissions associated with crop production, feed additive production, enteric methane, farm management, and manure storage were calculated and expressed as kg CO_2_ equivalents (CO_2_e) per kg of energy corrected milk. The total greenhouse gas emissions from baseline, two 3-nitroxypropanol and three nitrate scenarios were 1.12, 0.993, 0.991, 1.08, 1.07, and 1.09 kg CO_2_e/kg energy corrected milk. The average net reduction rates for 3-nitroxypropanol and nitrate were 11.7% and 3.95%, respectively. In both cases, using the feed additives on the whole herd slightly improved overall carbon footprint reduction compared to limiting its use during lactation phase. Although both 3-nitroxypropanol and nitrate had effects on decreasing the total greenhouse gas emission, the former was much more effective with no known safety issues in reducing the carbon footprint of dairy production in California.

## Introduction

The main greenhouse gases (GHG) emissions from agricultural food production include nitrous oxide (N_2_O), carbon dioxide (CO_2_) and methane (CH_4_). Livestock sector contributes to approximately 14.5% of global anthropogenic GHG emissions with 80% attributed to CH_4_ production from enteric fermentation and manure management from ruminants [1]. Dairy production is the third largest agricultural industry in the United States with total milk production increasing 13% over the past decade reaching over 215 billion pounds in 2019 [2]. California, as the top dairy production state accounted for over 20% of the total milk production with 1.73 million cows [2, 3].

The dairy industry has an environmental impact including GHG emissions related to crop production, enteric and manure CH_4_, water resource for feed production, excretion of nitrogen and phosphorus, and land management [4]. There are several mitigation strategies developed to reduce GHG emission from dairy, but most are modest in magnitude and some not applicable to California [5]. In the last few years, several feed additives have been developed to reduce enteric CH_4_ emissions with varying results. Two of the most studied feed additives that substantially reduce enteric CH_4_ emissions include 3-nitroxypropanol (3NOP) and nitrate (e.g., [6–9]). 3-nitroxypropanol (also known as Bovaer in Europe), is a synthetic compound that inhibit Methylcoenzyme M reductase, the enzyme that catalyzes the methane-forming step in the rumen [10]. Dijkstra et al. [8] conducted a meta-analysis to evaluate the anti-methanogenic effects of 3NOP and concluded that on average CH_4_ production (g day^−1^) was reduced 32.5% and CH_4_ yield (g kg dry matter intake^−1^) 29.3%. However, 3NOP appeared to be more effective in dairy reducing CH_4_ production 39.0% compared to 22.2% in beef. The authors also reported that the effectiveness of 3NOP was positively related to dose, but impaired by increased neutral detergent fibre (NDF) content in the diet.

Nitrate as an electron acceptor has been studied as dietary feed additive for ruminants to inhibit CH_4_ emission. Nitrate is reduced to nitrite and further reduced to ammonia, which is highly competitive with methanogens for hydrogen utilization in rumen due to a greater Gibbs energy changes than the CO_2_ to CH_4_ pathway [11]. The anti-methanogenic effect of nitrate have been investigated *in vivo* in various studies using beef steers, dairy cows, sheep, and goats [9, 12–15]. Van Gastelen et al. [16] conducted a meta-analysis and demonstrated that CH_4_ production consistently decreased when feeding nitrate to different ruminant animals. A recent meta-analysis by Feng et al. (unpublished) indicated that nitrate reduced CH_4_ production 14.4% and CH_4_ yield 11.4% in a dose-response manner. The main concern in using nitrate as a feed additive is the potential for nitrate toxicity. Nitrite as a result of nitrate reduction can accumulate in the animal and absorbed into blood. Increases in blood nitrite causes an increase in the concentration of methemoglobin, which can be fatal to animals. However, a denitrifying probiotic, *Paenibacillus fortis,* has been identifying as a way to enhance nitrite detoxification in nitrate treated ruminants [17]. The objective of this study was to estimate the net GHG emissions in California dairy system holistically based on supplementation of 3NOP and nitrate to the basal diet.

## Materials and methods

The study was based on a life cycle assessment (LCA) conducted for the dairy industry in California [4]. The feed ingredients used by Naranjo et al. [4] were adjusted and recalculated using NRC (2001). The impact of producing the feed additives 3NOP and nitrate was integrated in the LCA model. Energy corrected milk (ECM) was used as the functional unit and all emissions were calculated and standardized to 1 kg of ECM. The LCA conformed to Food and Agriculture Organization of the United Nations (FAO) Livestock Environmental Assessment Protocol guidelines.

The milk production supply chain in California from cradle to farm gate was considered the system boundary of the LCA including production of the feed additives. Specifically, these include: crop production, feed additives production, farm management, enteric methane, and manure storage (Fig 1). The system boundary considered emissions associated with on-farm activities, pre-farm production, and transportation of major productions up to the animal farm gate. Emissions for further activities after the products left the farm gate were not accounted in the system because they were considered to be treated in the same way for all scenarios.

**Fig 1.**
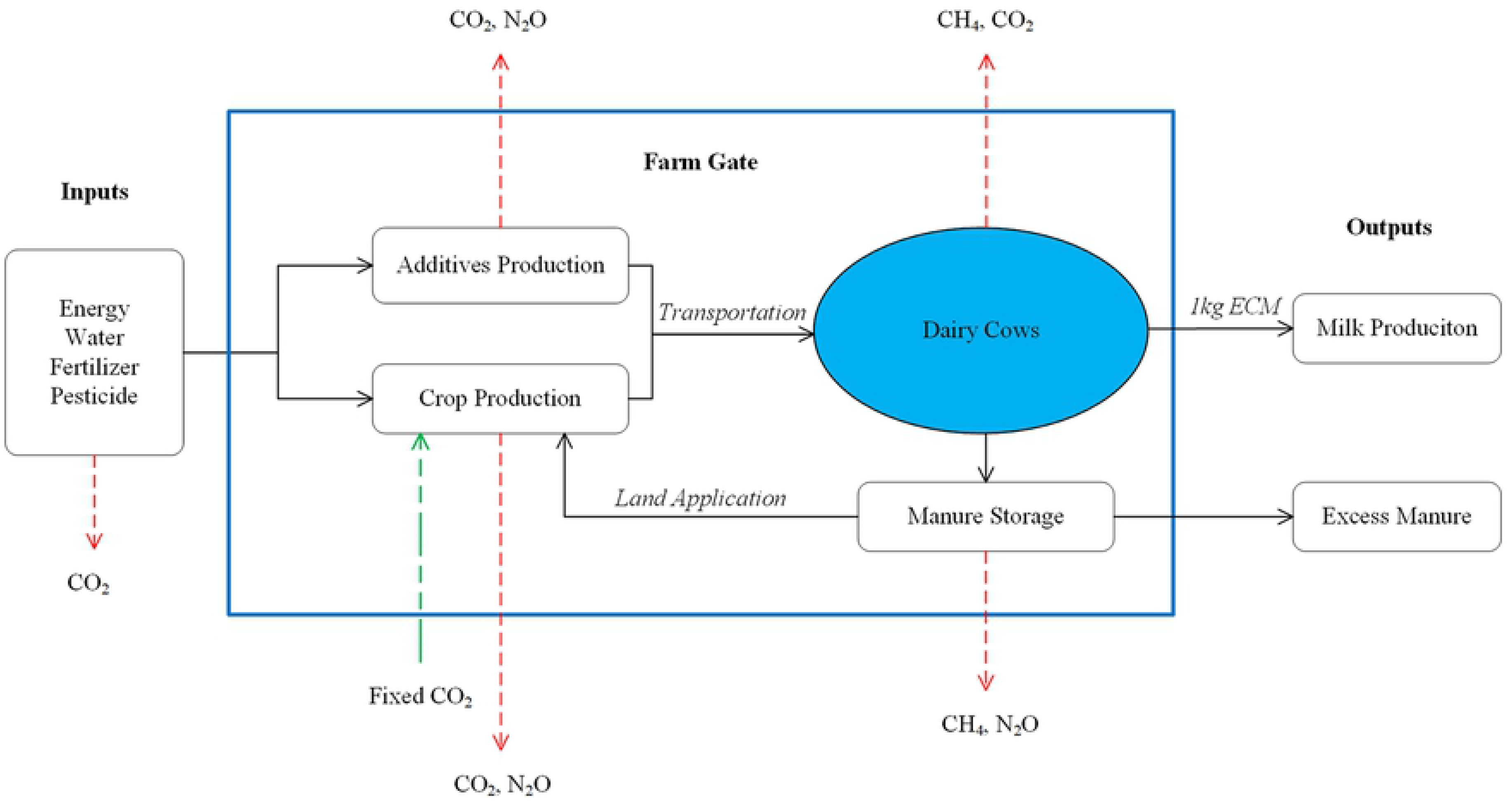
System boundary of the life cycle assessment for California milk production.

### Mitigation scenarios

Data were collected from USDA National Agricultural Statistical Service (USDA-NASS) and Economic Research Service (USDA-ERS), California Department of Food and Agriculture (CDFA), peer-reviewed literature and other published resources. The GHG emissions from each process in the LCA were estimated using the average conditions for dairy cattle in California as described by model 2 in Naranjo et al. [4]. The baseline scenario used California representative diets collected from CDFA reports. Average data from 2013 to 2015 represented the diets for year 2014 in the current analysis, which is the reference year. The diets for dairy cows at different growth stages including calf up to 1 year, heifer, pregnant heifer, close-up heifer, high lactating cow, and dry cow were weighted based on a whole production cycle. We assume 4 lactations to be the average life span of a California dairy cow. The crop production for baseline included the activities related to producing feed, and use of land, water, fertilizers, pesticides and herbicides. Additionally, energy used for machine operation, irrigation, and transportation was included. Data from USDA-NASS Quick Stats [18], USDA farm and ranch irrigation reports [19], California specific agricultural reports [20, 21], USDA-ERS reports [22], University of California crop cost and return studies [23], and values published in literatures [24] were used to estimate the emissions during the crop production. Enteric CH4 emissions, farm management, energy and water used for producing crop, feeding cattle, cooling livestock facilities, animals, and milk, sanitation, cleaning, and dealing with onsite waste were according to Naranjo et al. [4]. Similarly, manure methane and nitrous oxide (N_2_O) emissions were based on methodology described by Naranjo et al. [4].

Two scenarios were developed to estimate net mitigation effect of supplementing 3NOP to typical dairy diet in California. In scenario 1, all dairy cows were simulated to consume a diet that contains 3NOP only during lactation. In scenario 2, 3NOP was supplement to the diet at all growing stages within a life cycle. The basal diets were the same as in the baseline and 3NOP was supplemented at a rate of 127 mg/kg dry matter (DM) in both scenarios.

Nitrate as a non-protein nitrogen source for cattle is usually used to replace other non-protein N sources such as urea [25, 26]. Urea is not typically used as a nitrogen source in California representative diets, so nitrate was simulated to partially replace dietary protein in diets to keep similar N supply for all nitrate scenarios. In nitrate scenario 1, all dairy cows were simulated to consume a diet that contained nitrate only during lactation. Nitrate was supplemented to dairy cows at all stages in nitrate scenarios 2 and 3. In nitrate scenario 2, high protein meal (e.g. corn gluten, soybean meal, and distillers dried grain and solubles) was replaced by dietary nitrate on an equivalent nitrogen basis with no adjustment for dry matter intake (DMI). In nitrate scenario 3, DMI was adjusted using low protein meal (e.g. corn grain, and wheat silage) to the baseline levels after replacing high protein meal with nitrate additives. Nitrate was supplemented to dairy cattle at a rate of 16.7 g/kg of DM for all the 3 nitrate treatment scenarios.

### Emission associated with production and use of additives

#### 3-nitroxypropanol

The carbon footprint of emissions associated with 3NOP production were assumed to be 52 kg CO_2_e/kg 3NOP produced (DSM Nutritional Products, Ltd., pers. comm.). Moreover, with the improvement of process optimization, the carbon footprint of 3NOP could drop to 35 kg CO_2_e/kg 3NOP (DSM Nutritional Products, Ltd., pers. comm.). The total GHG emissions from 3NOP production were estimated using both factors and the results were reported as mean with standard error to evaluate the effect of 3NOP emission factors on net GHG emissions. The transportation of 3NOP was calculated based on shipping from the producer (DSM Nutritional Products, Ltd., registered in Ontario, CA) and dairy farms in California by truck. The average distance used to estimate the emissions related to 3NOP transportation was weighted according to the milk production amount in California counties in 2014 [27].

The magnitude of enteric CH_4_ emission reduction as a result of supplementing 3NOP was calculated based on an updated version of a meta-analysis conducted by Dijkstra et al. [8] on the anti-methanogenic effects of 3NOP. Four more recent references related to 3NOP effect on CH_4_ emissions were added to the previous analysis to extend the accuracy and robustness of the meta-analytical model. The updated database included treatment means from Martinez-Fernandez et al. [28] (beef; 1 treatment), Vyas et al. [29] (beef; 2 treatments), Kim et al. [30] (beef; 4 treatments), and van Wesemael et al. [31] (dairy; 2 treatments). The final mixed-effect models for CH_4_ production in the updated meta-analysis indicated effectiveness of 3NOP at mitigating CH_4_ production was positively associated with 3NOP dose, and negatively associated with NDF content. Similar to the previous study, supplementation of 3NOP had stronger anti-methanogenic effects in dairy cows compared to beef cattle, at a slightly greater magnitude of mitigation. The following equations were used to calculate the mitigation effect of 3NOP that includes dose, NDF content and either dairy (Eq 1) or beef (Eq 2):

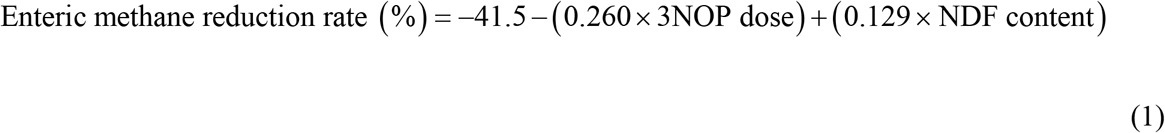

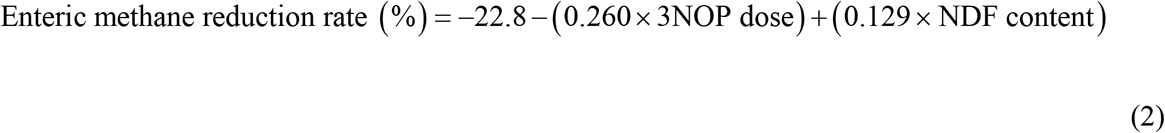

The equations were centered on the mean value of 127 mg 3NOP kg DM^−1^ and 326 g NDF kg DM^−1^. Therefore, the methane reduction rates were adjusted for each cattle type when the NDF content in the 3NOP supplemented scenarios varies from the default centered value. The NDF contents for different growing stages of dairy cows in California used in this study were calculated using NRC (2001) based on ingredients supplied (Table 1). In 3NOP scenario 1, enteric CH_4_ emitted from lactating cows was reduced 38.8%, which includes adjustment for NDF content (Table 1). In scenario 2, if the cows were not lactating, the emission reduction rate was assumed to be similar to beef cattle so Eq 2 was applied. The enteric CH_4_ reduction rates for various life-stages is given in Table 1.

**Table 1.** Enteric methane reduction rates and total emissions per life cycle at different dairy growing stages for baseline and 3NOP treatment scenarios.

| Cattle Stage | NDF<sup>a</sup><br>(g/kg DM) | Control |  | 3NOP 1 |  | 3NOP 2 |  |
| --- | --- | --- | --- | --- | --- | --- | --- |
|  |  | Reduction (%) | CH<sub>4</sub> emission (kg/lifetime) | Reduction (%) | CH<sub>4</sub> emission (kg/lifetime) | Reduction (%) | CH<sub>4</sub> emission (kg/lifetime) |
| Calf up to 1 year | 250 | NA | NA | NA | NA | NA | NA |
| Heifer | 419 | 0 | 10.6 | 0 | 10.6 | -11.1% | 9.4 |
| Pregnant heifer | 496 | 0 | 73.8 | 0 | 73.8 | -1.1% | 72.9 |
| Close up heifer | 425 | 0 | 9.2 | 0 | 9.2 | -10.3% | 8.2 |
| High lactating cow | 349 | 0 | 575.8 | -38.8% | 352.4 | -38.8% | 352.4 |
| Dry cow | 474 | 0 | 60.9 | 0 | 60.9 | -4.0% | 58.5 |

<sup>a</sup> NDF content in diets for baseline and 3NOP scenarios are same.

The GHG emissions from the farm management and manure management processes in the LCA for 3NOP scenarios were the same as for the baseline scenario because we assumed no residues and by-products from the 3NOP production process. Nkemka et al. [32] confirmed that there was no residual effect on anaerobic digestion of the manure from beef cattle fed diets supplemented with 3NOP.

#### Nitrate

Nitrate was assumed to be supplemented to dairy diets as Calcium nitrate (Ca(NO_3_)_2_). Brentrup et al. [33] reported carbon footprint associated with Ca(NO_3_)_2_ production were estimated to be 1.76 kg CO_2_e kg Ca(NO_3_)_2_^−1^ in USA and 0.67 kg CO_2_e kg Ca(NO_3_)_2_^−1^ produced in Europe. Total emissions associated with Ca(NO_3_)_2_ production were calculated using both carbon footprint values for USA and Europe, and the emissions from nitrate production process are reported as the mean with standard deviation. Emissions related to transportation of Ca(NO_3_)_2_ were calculated based on the shipping distance between supplier and dairy farms in California. Several chemical companies supply Ca(NO_3_)_2_ within California and the plant with the minimum travel distance (by truck) to each county was assumed as its Ca(NO_3_)_2_ supplier. The overall average distance was weighted based on the milk production in California counties in 2014 [27] and used for emission calculations related to chemical transportations. Feed production for different nitrate treatment scenarios were recalculated based on the replacement of high protein meals by dietary nitrate to provide equivalent nitrogen compared to the diets for control scenario at each growing stage using NRC (2001).

The anti-methanogenic effects of nitrate were calculated based on equations developed by Feng et al. (unpublished). Meta-analytical results indicated nitrate effect on enteric CH_4_ production to be significantly affected by nitrate dose, cattle type, and DMI and the mitigation effects of nitrate on CH_4_ production was greater in dairy cows compared to beef cattle. The reduction rates for enteric CH_4_ emissions estimated by the meta-analytical model were given in Eq 3 for dairy cattle and Eq 4 for beef cattle.

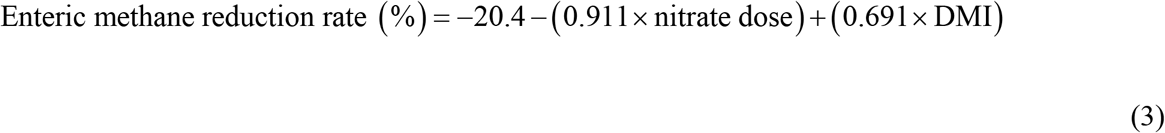

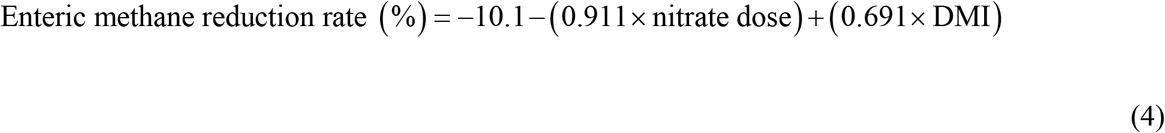

The equations are centered on mean nitrate dose and mean DMI of the database, which were 16.7 g kg of DM^−1^ and 11.1 kg day^−1^, respectively. We kept the average as the dose of nitrate supplementation in the scenarios evaluated in this study. The DMI for different growing stages for baseline in this study were estimated from CDFA reports to represent the daily feed intakes of dairy cows in California (Table 2). In nitrate scenario 1, DMI for high lactating cow slightly dropped from baseline of 22.6 kg day^−1^ to 22.3 kg day^−1^ due to the replacement of high protein meal by concentrate nitrate and enteric CH_4_ emitted was reduced 13.6% when adjusted for DMI (Table 2). In nitrate scenario 2, high protein ingredients were replaced by nitrate for all growing stages which resulted in DMI differences compared to the baseline. The reduction rates of enteric CH_4_ emissions were estimated by Eq 4 for beef cattle, which were 14.7%, 11.4%, 10.7%, 13.6%, and 10.3% for heifer, pregnant heifer, close up heifer, high lactating cow, and dry cow, respectively (Table 2). In nitrate scenario 3, DMI were adjusted back to the baseline levels and the emission reduction rates were calculated using the same approach as nitrate scenario 2. The enteric CH4 emissions for heifer, pregnant heifer, close up heifer, high lactating cow, and dry cow were reduced by 14.7%, 11.0%, 10.3%, 13.4%, and 10.0%, respectively (Table 2).

**Table 2.** Enteric methane reduction rates and total emissions per life cycle at different dairy growing stages for control and nitrate treatment scenarios.

| Cattle Stage | Control |  |  | Nitrate 1 |  | Nitrate 2 |  | Nitrate 3 |  |
| --- | --- | --- | --- | --- | --- | --- | --- | --- | --- |
|  | DMI<sup>a</sup> (kg/d) | Reducti on (%) | CH<sub>4</sub> emission (kg/lifetime) | Reducti on (%) | CH<sub>4</sub> emission (kg/lifetime) | Reducti on (%) | CH<sub>4</sub> emission (kg/lifetime) | Reducti on (%) | CH<sub>4</sub> emission (kg/lifetime) |
| Calf up to 1 year | 4.1 | NA | NA | NA | NA | NA | NA | NA | NA |
| Heifer | 5.8 | 0 | 10.6 | 0 | 10.6 | -14.7% | 9.7 | -14.7% | 9.8 |
| Pregnant heifer | 11.1 | 0 | 73.8 | 0 | 73.8 | -11.4% | 65.4 | -11.0% | 69.5 |
| Close up heifer | 12.1 | 0 | 9.2 | 0 | 9.2 | -10.7% | 8.0 | -10.3% | 8.2 |
| High lactating cow | 22.6 | 0 | 575.8 | -13.6% | 493.1 | -13.6% | 493.1 | -13.4% | 500.6 |
| Dry cow | 12.6 | 0 | 60.9 | 0 | 60.9 | -10.3% | 53.3 | -10.0% | 55.4 |

<sup>a</sup> DMI for control and Nitrate scenario 3 are same. DMI for Nitrate scenario 1 are same as for control except for high lactating cows (DMI = 22.3 kg/d). DMI for Nitrate scenario 2 in different cattle stages are 4.1 (Calf up to 1), 5.8 (Heifer), 10.5 (Pregnant heifer), 11.6 (Close up heifer), 22.3 (High lactating cow), and 12.1 (Dry cow), respectively.

We assumed there were no residues and by-products from nitrate production and the total GHG emissions from farm management process for nitrate treatment scenarios including on-farm energy and water usage were not affected by nitrate additives. Methane emissions from manure storage were calculated as a function of volatile solids excreted [34], which was associated with NDF content, crude protein content and DMI [35]. As the dietary ingredients and DMI for nitrate scenarios varied with the adjustment of nitrate additives, the total GHG emissions from manure management were recalculated based on the different nitrate feeding scenarios.

## Results and discussion

### 3-nitroxypropanol

The GHG emissions from crop production, farm management, enteric CH4 and manure storage for baseline were 0.174, 0.0608, 0.432, and 0.457 kg CO_2_e kg of ECM^−1^ produced in California, respectively (Fig 2). Total GHG emissions from crop production, farm management, and manure storage were not affected by feeding 3NOP to dairy cows. There was no significant effect of 3NOP on DMI in cattle (e.g., [36, 37]), therefore, the total amount of basal diets consumed were assumed to be similar in cows supplemented with or without 3NOP and GHG emissions from feed production remained the same. The mean GHG emissions related to production of 3NOP in scenario 1 was 3.23 g CO_2_e kg ECM^−1^ which was lower than 3.92 g CO_2_e kg ECM^−1^ in scenario 2 because 3NOP was only fed to lactating cows in scenario 1. Enteric CH_4_ emissions were 0.298 and 0.295 kg CO_2_e kg ECM^−1^ for 3NOP scenarios 1 and 2, respectively, which were reduced 31.0% and 31.7% compared to baseline, respectively, due to the inhibition effect of 3NOP on CH_4_ production. Accounting for emissions from 3NOP production, the net enteric methane emission reduction was 30.3% in scenario 1 and 30.8% in scenario 2.

**Fig 2.**
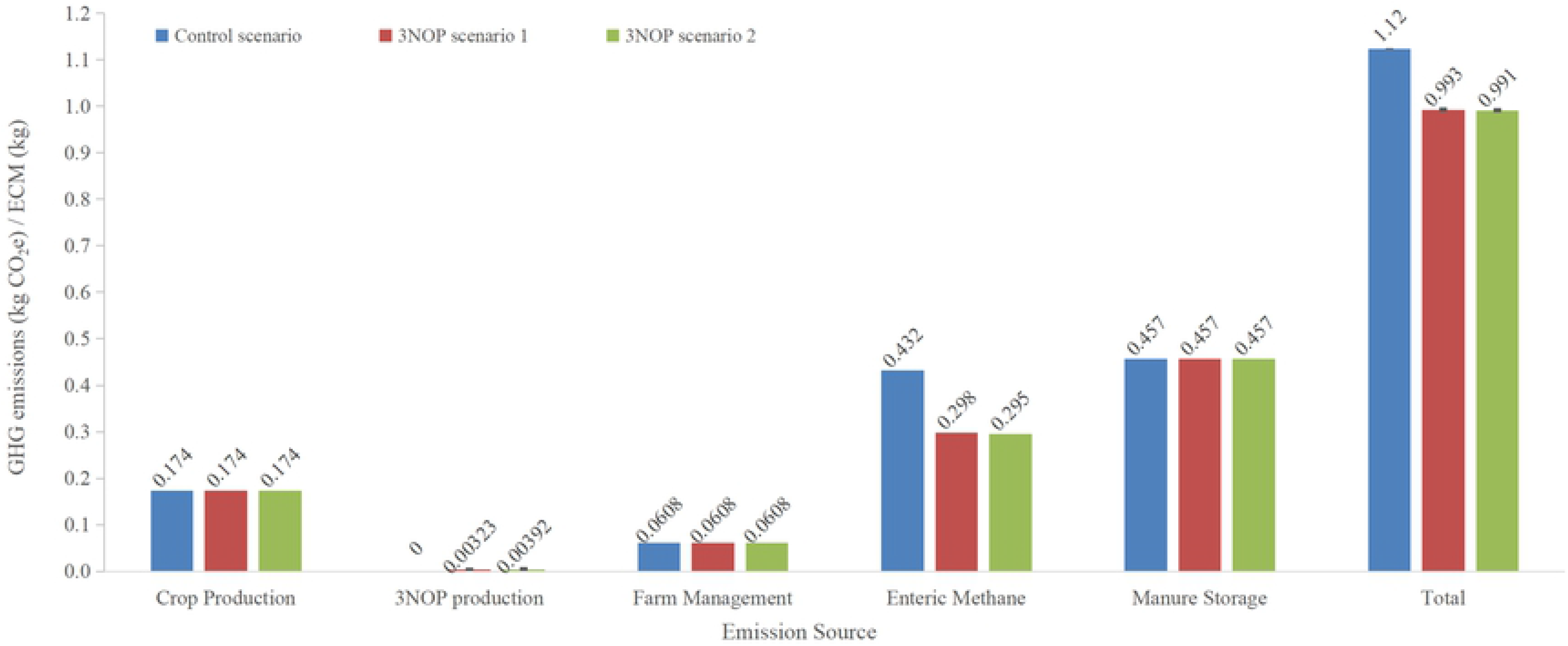
Comparison of global warming potential (GWP) by emission source for control and 3-nitroxypropanol (3NOP) scenarios 1 and 2 in California dairy cows.

The total GHG emissions for baseline and 3NOP treatment scenarios 1 and 2 were 1.12, 0.993 and 0.991 kg CO_2_e kg ECM^−1^, respectively (Fig 2). Feeding 3NOP to dairy cows resulted in a net reduction of total GHG emission of 11.6% in 3NOP scenario 1 and 11.8% in 3NOP scenario 2 compared to the baseline. Using 3NOP for dairy cows at all growing stages only further reduced 0.2 percentage points more compared to limiting 3NOP supplementation during lactation. The small change in mitigation effect of GHG emissions in 3NOP scenario 2 compared to scenario 1 was due to less effectiveness of 3NOP on non-lactating cattle and relatively shorter period spent in non-productive phase (Table 1). The GHG emissions estimated from groups supplemented with 3NOP at 86 mg kg DM^−1^ based on two Canadian dairy farms were 1.03 (fed 3NOP during lactations only) and 0.98 kg (fed 3NOP to entire herd) CO_2_e kg ECM^−1^ which was a reduction of 14.9% and 19.0% compared to their baseline, respectively [38]. The study also investigated the GHG emissions from two dairy farms in Australia and the reductions in GHG emissions compared to their baseline were 14.4 to 14.7%, when 3NOP was fed for 300 days of lactation (86 mg/kg DM). The net reductions of GHG emissions in California dairy farms were lower than estimated by Alvarez-Hess et al. [38] mainly because in California, CH_4_ from manure management is greater compared to Canadian and Australian conditions, therefore, the overall effect of 3NOP was slightly diluted.

The GHG emissions associated with 3NOP production for scenarios 1 and 2 were 3.86 (3NOP scenario 1) and 4.69 (3NOP scenario 2) g CO_2_e kg ECM^−1^, respectively, assuming 3NOP carbon footprint of 52 kg CO_2_e kg^−1^ and 2.60 (3NOP scenario 1) and 3.16 (3NOP scenario 2) g CO_2_e kg ECM^−1^, respectively, using manufacturer reported values of 35 kg CO_2_e kg 3NOP^−1^. This indicates that with the improvement of manufacturing process, the GHG emissions from 3NOP production can be reduced 32.6% and improve net impact of 3NOP in reducing enteric emissions.

### Nitrate

The total GHG emissions and estimates of the various components in dairy cattle supplemented with nitrate is given in Fig 3. In nitrate scenario 1, the mean GHG emissions associated with nitrate production was 0.0169 kg CO_2_e kg ECM^−1^ and 0.0203 kg CO_2_e kg ECM^−1^ in nitrate scenarios 2 and 3 due to differences in the phases of dairy production that nitrate was included. The differences in carbon footprint of Ca(NO_3_)_2_ production was mainly due to a catalyst technology developed in Europe [33]. The GHG emissions calculated with carbon footprint value for Ca(NO_3_)_2_ in USA (1.76 kg CO_2_e kg per Ca(NO_3_)2^−1^ produced) were 0.0237 kg CO_2_e kg ECM^−1^ for nitrate scenario 1, and 0.0285 kg CO_2_e kg ECM^−1^ for nitrate scenarios 2 and 3. Using the European carbon footprint (0.67 kg CO_2_e kg Ca(NO_3_)2^−1^ produced), the GHG emissions from nitrate production was 0.0101 kg CO2e kg ECM^−1^ for nitrate scenario 1, and 0.0122 kg CO2e kg ECM^−1^ for nitrate scenarios 2 and 3. The GHG emissions from nitrate production (averaged from three scenarios) decreased 57.3% based on European values compared to those in USA.

**Fig 3.**
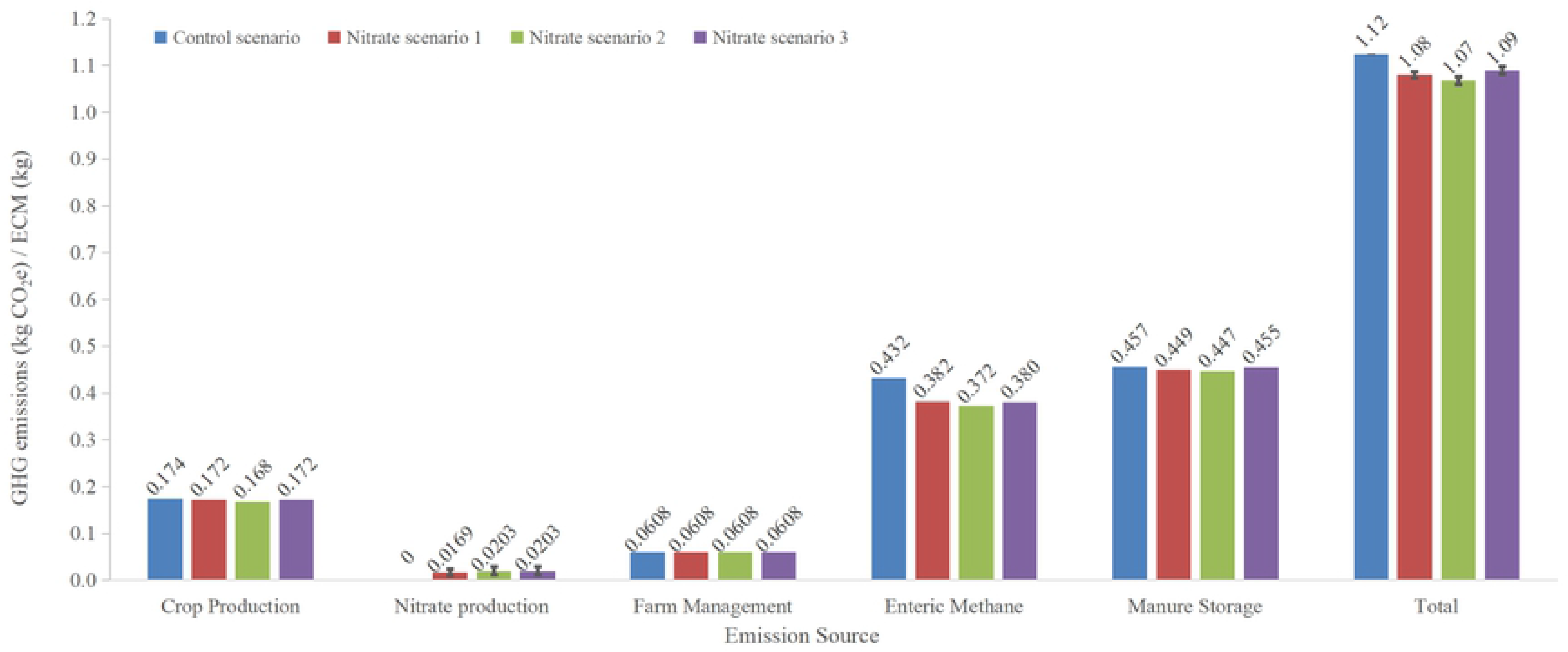
Comparison of global warming potential (GWP) by emission source for control and nitrate scenarios of dairy cows in California.

The GHG emissions related to crop production was 0.174 kg CO_2_e kg ECM^−1^ for the baseline, and reduced to 0.172, 0.168, and 0.172 CO_2_e kg ECM^−1^ for nitrate scenarios 1, 2, and 3, respectively, which was mainly caused by the decline in the amount of protein that was replaced by nitrate. Although replacing high protein sources such as corn gluten, and soybean meals reduced the GHG emissions of feed production, emissions from nitrate production were much greater in comparison. The DMI for scenario 3 was adjusted back to baseline level, therefore, the GHG emissions from crop production in nitrate scenario 3 was 0.004 CO_2_e kg ECM^−1^ greater than in scenario 2. The GHG emissions from manure storage were 0.457, 0.449, 0.447, 0.455 kg CO_2_e kg ECM^−1^ in control and three nitrate scenarios, respectively. The differences of GHG emissions from manure management among nitrate scenarios were associated with the variations in dietary NDF content, crude protein content, and DMI of adjusted diets. Enteric CH_4_ emissions from nitrate scenarios 1 to 3 were 0.382, 0.372, and 0.380 kg CO_2_e kg ECM^−1^, respectively, which were reduced by 11.5%, 13.9%, and 12.0% respectively, compared to CH_4_ emissions from control scenario (0.432 CO_2_e kg ECM^−1^) based on values calculated for CH_4_-mitigating effect of dietary nitrate (Table 2). The net reduction in enteric methane emission (and GHG from nitrate production) was calculated to be 7.58, 10.4 and 8.42% for nitrate scenarios 1 to 3, respectively. The GHG emissions from farm management were the same for control and all nitrate scenarios which was 0.0608 kg CO_2_e kg ECM^−1^ (Fig 3).

The total GHG emissions for control scenario was 1.12 kg CO_2_e kg ECM^−1^, while with supplementing dietary nitrate to dairy cows in California, the total GHG emissions were 1.08, 1.07, and 1.09 kg CO_2_e kg ECM^−1^ (Fig 3), respectively, in nitrate scenarios 1, 2, and 3. The total GHG emissions for three nitrate scenarios were reduced 3.82%, 4.96%, and 3.07% compared to the control scenario, respectively. In scenario 2, nitrate was fed to all growing stages with the largest amount of replaced basal diet resulting in the lowest GHG emissions from crop production and enteric CH_4_. Therefore, scenario 2 showed the greatest net reduction of GHG emissions, which was reduced 1.15% and 1.89% more compared to scenarios 1 and 3, respectively. These results were lower than the reduction rates estimated by Alvarez-Hess et al. [38] who reported that the GHG emissions went down from 1.21 kg CO_2_e kg ECM^−1^ (baseline) to 1.13 (a reduction of 6.61%) and 1.10 (a reduction of 9.09%) kg CO_2_e kg ECM^−1^ when nitrate was fed at a rate of 21 g/kg DM to lactating cows only and to the entire herd, respectively. Alvarez-Hess et al. [38] used an average reduction rate of 23% for CH_4_ emissions with supplementing nitrate, while we assumed enteric CH_4_ emissions were decreased 20.4% for lactating cows and 10.1% for dairy cows under other growing stages without affecting milk production when the nitrate was fed at a rate of 16.7 g kg of DM^−1^ and the DMI of 11.1 kg day^−1^. This may explain the relatively lower net reduction of total GHG emissions in the current study.

### Comparison of 3-nitroxypropanol and nitrate additives

Total GHG emissions from baseline (1.12 kg CO_2_e kg ECM^−1^) were lower than values published in several previous studies. For example, Gerber et al. [39] reported the GHG emissions in North America to be 1.20 kg CO_2_e kg ECM^−1^ and Thoma et al. [40] reported 1.23 kg CO_2_e kg ECM^−1^. In Canada, Guyader et al. [41] reported the GHG emissions varied from 1.18 to 1.52 kg CO_2_e kg ECM^−1^ for a dairy farm and Alvarez-Hess et al. [38] reported 1.21 kg CO_2_e kg ECM^−1^, but in two Australian dairy farms the authors reported 1.09 and 0.97 kg CO_2_e kg ECM^−1^, respectively, which were lower than the value estimated in the present study. Enteric CH4 and manure management are the major sources of GHG emissions in the dairy sector [40]. Emissions from manure storage accounted for 40.6% to 46.1% of the total GHG emissions, which contributed the largest amount to total GHG emissions in all scenarios. Enteric CH4 emissions from baseline scenario accounted for 38.4% of the total GHG emissions but the proportions of enteric CH_4_ emissions dropped to between 29.8% (3NOP scenario 2) to 35.4% (nitrate scenario 1). Crop production emitted 15.5% to 17.6% of total GHG emissions and the significant decrease in enteric CH_4_ emissions resulted in a proportional increase of GHG emissions of crop production in 3NOP scenarios. Only 0.3% to 1.9% of emissions were attributed to feed additives production in supplemental scenarios. The GHG emissions associated with farm management were the same for all scenarios.

Although both 3NOP and nitrate additives decreased the total GHG emissions, the mitigating effect of 3NOP was greater than nitrate reaching a highest reduction rate of 11.8% (3NOP scenario 2). The average net reduction rate of GHG emissions for 3NOP scenarios was 11.7% and supplementing 3NOP to dairy cows only during lactations or to the entire growing herds had a minor difference in the total GHG emissions. The mean net reduction rate of GHG emissions in dairy cows feeding nitrate was 3.95%. The greatest net GHG emissions achieved with nitrate was 4.96% with supplementation of nitrate to dairy cows in all growing stages. In addition to differences in effectiveness, nitrate is fed at an average rate of 16.7 g kg DM^−1^ compared to an average 3NOP rate of 127 mg kg DM^−1^ in this study. Therefore, much higher quantities of nitrate are required for enteric CH_4_ mitigation resulting in about 5.4 times GHG emission from production of the additive alone. Nitrate toxicity caused by the high methemoglobin levels in ruminants fed in greater quantities is a concern and currently not recommended as CH4 mitigating feed additives to cattle [42, 43].

## Conclusions

This study was conducted based on dairy cows in California and evaluated the mitigation effect of two effective feed additives—3NOP and nitrate, on total GHG emissions. The average net reduction rate of supplementing 3NOP and nitrate were 11.7% and 3.95%, respectively. 3NOP had a greater effect than nitrate on reducing total GHG emissions with a highest performance of 11.8%. Feeding 3NOP to only lactating cows or to the entire growth stages did not make significant difference in total GHG emissions. Considering California milk production of 18 billion kg in 2017, using nitrate on California dairy cows would reduce GHG emissions 0.90 billion CO_2_e and 3NOP 2.33 billion CO_2_e annually.

